# Healthy carriage of *Salmonella* within cattle lymph nodes is a key source of ground beef contamination with strains of clinical significance

**DOI:** 10.1101/2024.04.16.589703

**Authors:** Enrique Jesús Delgado-Suárez, Abril Viridiana García-Meneses, Elfrego Adrián Ponce-Hernández, Cindy Fabiola Hernández-Pérez, María Salud Rubio-Lozano, Nayarit Emérita Ballesteros-Nova, Orbelín Soberanis-Ramos

## Abstract

This study assessed the genetic relatedness, evolutionary dynamics, and virulence profile of *Salmonella* isolated from lymph nodes and ground beef from apparently healthy cattle over two years. For this purpose, we used a set of isolates of nine different serovars: Anatum (n=23), Reading (n=22), Typhimurium (n=10), London (n=9), Kentucky (n=6), Fresno (n=4), Give, Muenster, and monophasic 1,4,[5],12:i- (n=1 each). These isolates were subjected to whole genome sequencing, and assembled and annotated genomes were used for downstream bioinformatic analyses. Although lymph nodes and ground beef were collected and analyzed separately, we still observed clonality between isolates from both sources. This finding suggests that some *Salmonella* circulating in the gut may reach the lymphatic system, creating a dual reservoir for ground beef contamination. We also found evidence of *Salmonella* persistence across cattle cohorts, as we observed clonality between isolates collected in different years. Variation in the virulence and pathogenicity island profiles was limited, with minor differences that could not be associated with attenuated virulence in the bovine host. Conversely, isolates of all serovars, except Fresno, were genetically close to strains involved in human salmonellosis in different countries, highlighting the risk to public health posed by strains associated with the carrier state in cattle. Further research is needed to reveal the mechanisms by which *Salmonella* causes subclinical infections in cattle and persists for long periods within farm environments.

## Introduction

Ground beef is considered a relevant vehicle of human exposure to *Salmonella enterica* (from now on referred to as *Salmonella*) [1, 2], one of the leading causes of foodborne diseases globally [3]. Several *Salmonella* strains use certain niches in cattle (e.g. intestines and lymph nodes) [4-6] without causing any sign of illness, and through mechanisms that have not been elucidated. As the pathogen goes undetected, producers do not apply any control measures, which has been suggested as a plausible mechanism behind the emergence or predominance of certain *Salmonella* serovars in cattle [7]. Research in this area is crucial as preventing asymptomatic *Salmonella* infections in cattle has been ineffective.

Increasing evidence has shown that lymph nodes are a significant source of ground beef *Salmonella* contamination [8-10]. However, despite intensive research, little is known about how the pathogen reaches and survives within the lymph nodes of healthy animals. Nonetheless, certain *Salmonella* serovars have been shown to predominate in this niche (e.g. Anatum, Reading, Montevideo, Kentucky) [4, 9, 11], suggesting that they may be better adapted to lymph nodes. This phenomenon could be associated with differences in the pathogen’s gene repertoire, a topic that has been poorly studied, particularly in terms of the virulence profile. Moreover, apart from typing approaches based on serotyping and pulsed-field gel electrophoresis (PFGE) [10, 12], few studies have addressed the genetic relatedness and evolutionary dynamics within these *Salmonella* populations.

In this investigation, we used comparative genomics and phylogenetic analyses to study the epidemiology, virulence profile, and genetic relatedness of a set of *Salmonella* isolates previously collected by our research team over a 2-year period [9], from cattle lymph nodes and ground beef.

## Materials and methods

Animal Care and Use Committee approval was not obtained for this study since live animals were not directly involved in the experiment. In this study, we used 77 *Salmonella* isolates of nine different serovars (Anatum, n=23; Reading, n=22; Typhimurium, n=10; London, n=9; Kentucky, n=6; Fresno, n=4; and Give, Muenster, and monophasic 1,4,[5],12:i- (n=1 each). These isolates were collected in 2017-2018 from 1,545 samples of cattle lymph nodes and ground beef, which were obtained from a carcass retailer located in Mexico City, as reported in a previous study by our research team [9]. Since we aimed to assess the genetic relatedness of isolates circulating in lymph nodes and ground beef, both sample types were collected and analyzed separately [9].

### Whole-genome sequencing and quality control of raw reads

Fresh colonies were grown overnight in trypticase soy broth (TSB) at 37 ºC under agitation. Next, 1 mL of the TSB was centrifuged at 5,000 x g for 10 min to obtain a cell pellet. Subsequently, the cell pellet was used to extract genomic DNA (gDNA) using the High Pure PCR Template Preparation Kit (Roche Molecular Systems, Inc., Switzerland). Afterwards, we quantified gDNA using a Qubit 3.0 fluorometer to ensure a minimum input of 1 ng of gDNA for the DNA library preparation workflow.

DNA libraries were prepared with the Nextera XT version 3 kit, and the sequencing was performed in an Illumina NextSeq platform (paired end 2 x 150 bp insert size, and a minimum estimated depth of coverage of 30x). Raw reads were uploaded to the National Center for Biotechnology Information (NCBI) repository and are publicly available through the accession numbers provided in this paper (Fig 1 and supplementary S1 Table).

**Fig 1.**
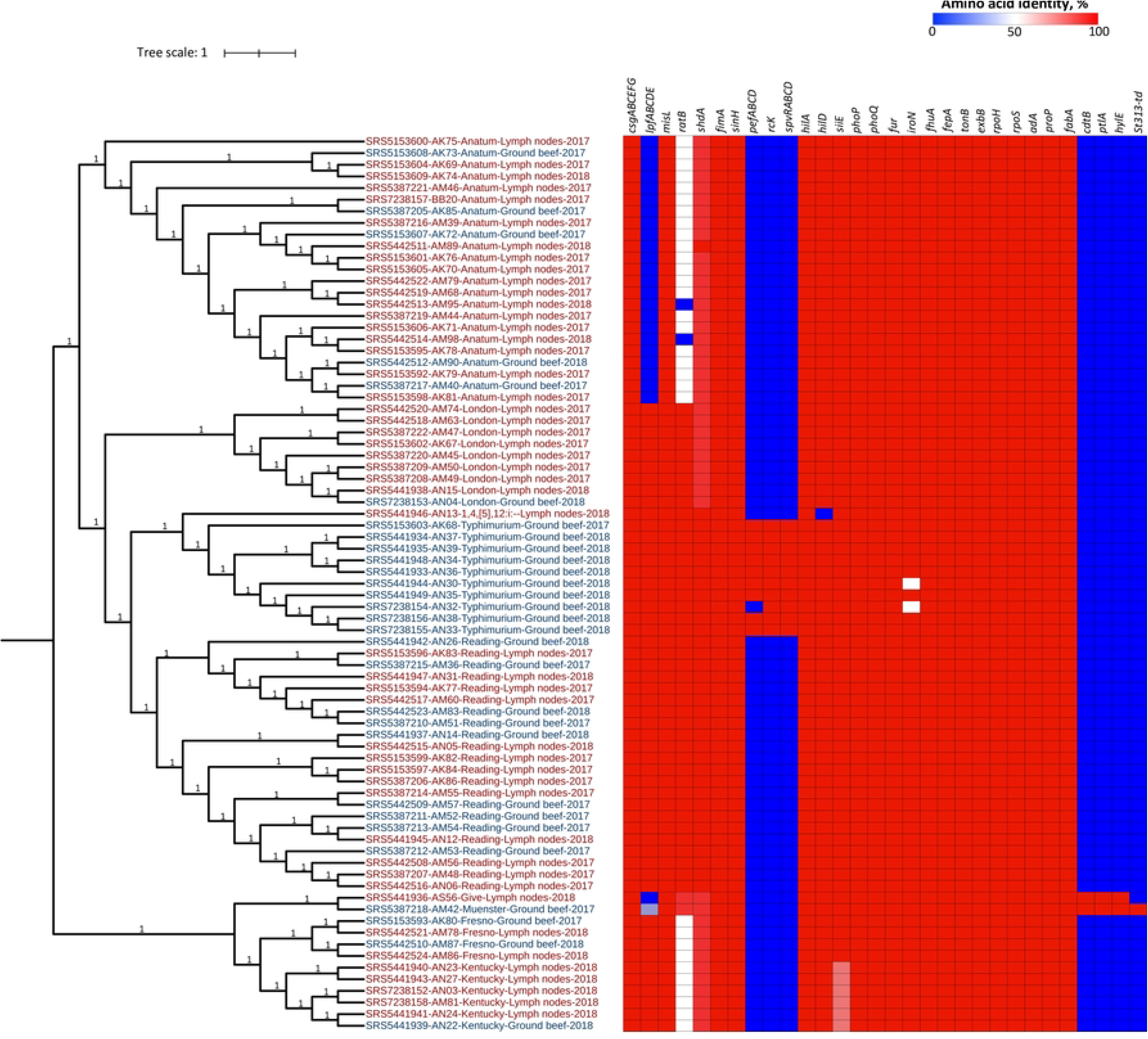
ML tree based on SNP analysis of 77 *Salmonella* isolates from cattle lymph nodes and ground beef. Tip labels are color coded according to isolation source and indicate NCBI accession, strain name, serovar, isolation source, and isolation year. The virulence profile is mapped onto the tree. The bootstrap values supporting each clade are indicated in the branches.

The quality of raw sequences was first assessed using FastQC [13]. Next, we used Trimmomatic version 0.39 [14] to remove Illumina adaptors and filter raw reads according to quality criteria. Trimmed sequences were analyzed again with FastQC to ensure that only high-quality reads (i.e. Q≥30) were used for genome assembly and downstream bioinformatic analyses.

### Genome assembly and annotation

We used SPAdes version 3.1.3.1 [15] to perform *de novo* genome assembly using trimmed sequences, while assembly quality was assessed using QUAST version 5.02 [16]. Data of assembly quality attributes was reported in our previous paper [17]. Genome annotation was conducted at the RAST web server using the RASTtk algorithm [18].

### Phylogenetic and evolutionary analysis

We conducted a maximum likelihood (ML) phylogenetic analysis based on single-nucleotide polymorphisms (SNP) using assembled genomes. For this purpose, SNPs were located, filtered, and validated for the 77 genomes using CSI Phylogeny version 1.4 [19], with default values. We used the *Salmonella* Typhi CT18 genome as a reference (accession GCA_000195995.1), although it was not included in the final phylogeny.

The resulting alignment was analyzed with RAxML version 8.0 [20] to generate an ML tree under the GTR+Γ model of nucleotide evolution at the CIPRES Science Gateway server version 3.3, with default values [21]. The resulting tree was edited using iTOL version 6.9 [22].

To assess the evolutionary dynamics of *Salmonella* circulating in lymph nodes and ground beef, we used the resulting phylogenetic tree to reconstruct the character state (isolation source) at ancestral nodes using Mesquite software, version 3.70, with a parsimony unordered model [23].

### Detection of virulence genes and Salmonella pathogenicity islands

We screened annotated genomes for the presence of major *Salmonella* virulence factors using the set of genes reported in the virulence factors database [24]. For this purpose, we performed a basic local alignment search tool (BLAST) analysis with the annotated genomes within RAST using the amino acid sequence of each reference protein and an e-value threshold of 10^-30^. We then used the resulting amino acid identity percentage to build a heatmap of the virulence profile using MORPHEUS software (https://software.broadinstitute.org/morpheus). The obtained virulence profile was mapped onto the ML tree mentioned above to identify whether it was associated with the isolation source of the isolates.

To identify the *Salmonella* pathogenicity islands (SPI) carried in the genomes, we first collected the reference sequences of SPIs 1-12 from the Pathogenicity Island Database [25]. Next, we conducted a BLAST atlas analysis using our study genomes and the concatenated nucleotide sequences of the 12 SPIs on the GView web server [26], with default values.

Finally, to investigate the risk posed by these *Salmonella* strains to public health, we analyzed the SNP clusters corresponding to each of our study isolates within the NCBI Pathogen Detection website (https://www.ncbi.nlm.nih.gov/pathogens). This analysis helped to assess the genetic proximity between our study isolates and those involved in human salmonellosis. The full list of SNP clusters analyzed in this study is provided in supplementary S1 Table.

## Results

### Phylogenetic and evolutionary analysis

The *Salmonella* population structure was relatively diverse. The SNP phylogeny divided the isolates into two divergent clades and seven SNP clusters (Fig 1). Nearly all isolates clustered according to serovar, except for the singletons of the serovars Muenster and Give, which were very close to each other. Moreover, we observed clonality (100 % bootstrap support) between some isolates of the same serovar across clusters, regardless of their isolation source. There was also clonality between the strains of most serovars, except Kentucky, despite being isolated in different years.

The reconstruction of character states at ancestral nodes showed that lymph nodes were the most parsimonious isolation source of the ancestors of all study isolates (Fig 2). However, the character state of some ground beef isolates remained unchanged across up to six lineage splitting events (i.e. isolates of serovar Typhimurium).

**Fig 2.**
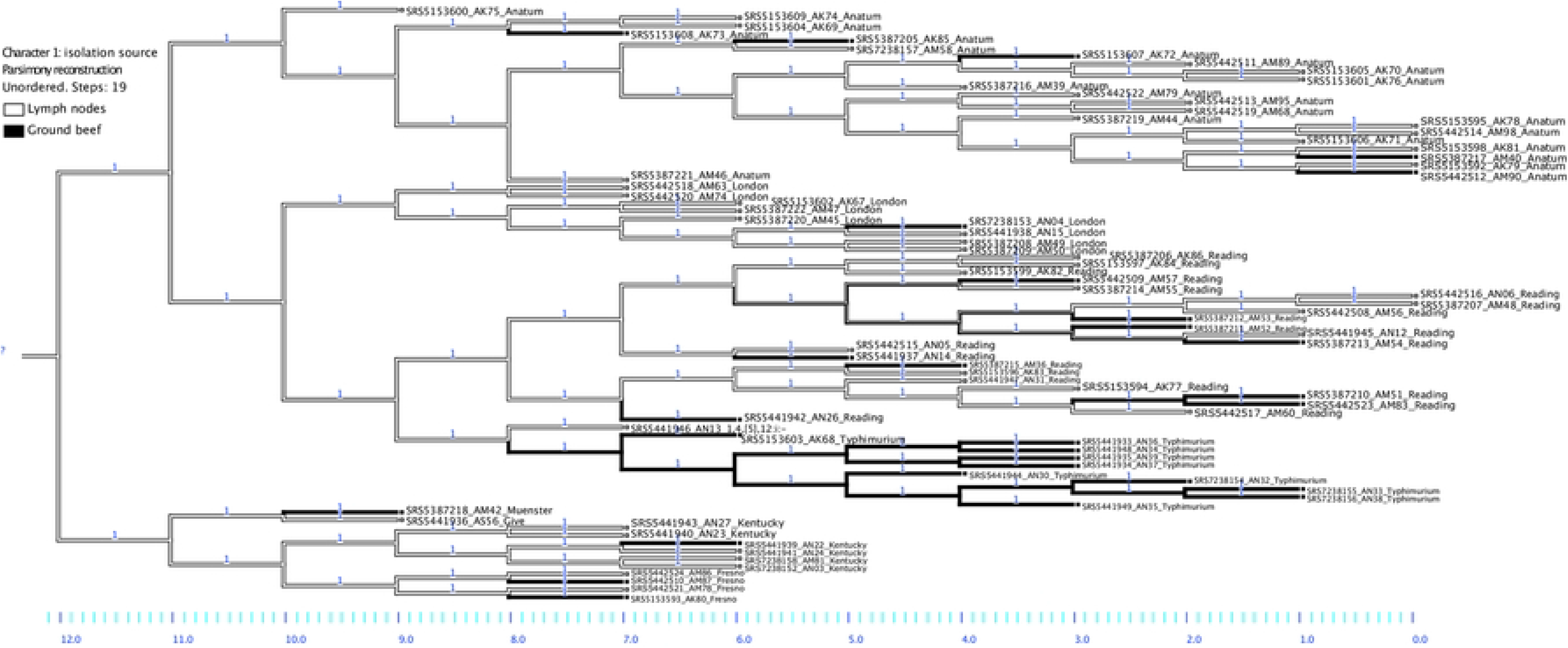
Reconstruction of the character state (isolation source) at ancestral nodes in a phylogenetic tree of 77 *Salmonella* isolates recovered from cattle lymph nodes (gray branches) and ground beef (black branches) in 2017-2018. NCBI accession, strain name, and isolation source of each isolate are indicated on the tip labels.

### Virulence and pathogenicity islands

Variations in virulence factors occurred mostly in genes encoding adhesion, toxins, and *Salmonella* virulence plasmid proteins (Fig 1). For instance, nearly one-third of the isolates lacked the long polar fimbriae operon (*lpfABCDE*). Moreover, isolates of serovar Anatum carried *ratA*, which is a variant with 47% sequence identity to *ratB* (a non-fimbrial adhesion factor), whereas two of them lacked this gene. In addition, *Salmonella* virulence plasmid genes (*pefABCD, spvRABCD, rcK*) were detected solely in serovar Typhimurium isolates, whereas only two isolates (serovars Give and Muenster) carried toxin-encoding genes (*cdtB, ptlA, hlyE*). Interestingly, the Muenster isolate carried the *st313-td* gene, which has been reported previously in strains of serovars Typhimurium and Dublin exhibiting invasive phenotypes in human infections [27].

Regarding pathogenicity islands, their sequences were uniform within the serovars. Therefore, we reported the results per serovar using a representative isolate of each serovar (Fig 3). In general, however, the SPI profile was similar across serovars as well. Nearly all isolates carried SPIs 1-6, 9, 11-12, and lacked SPIs 7, 8, and 10.

**Fig 3.**
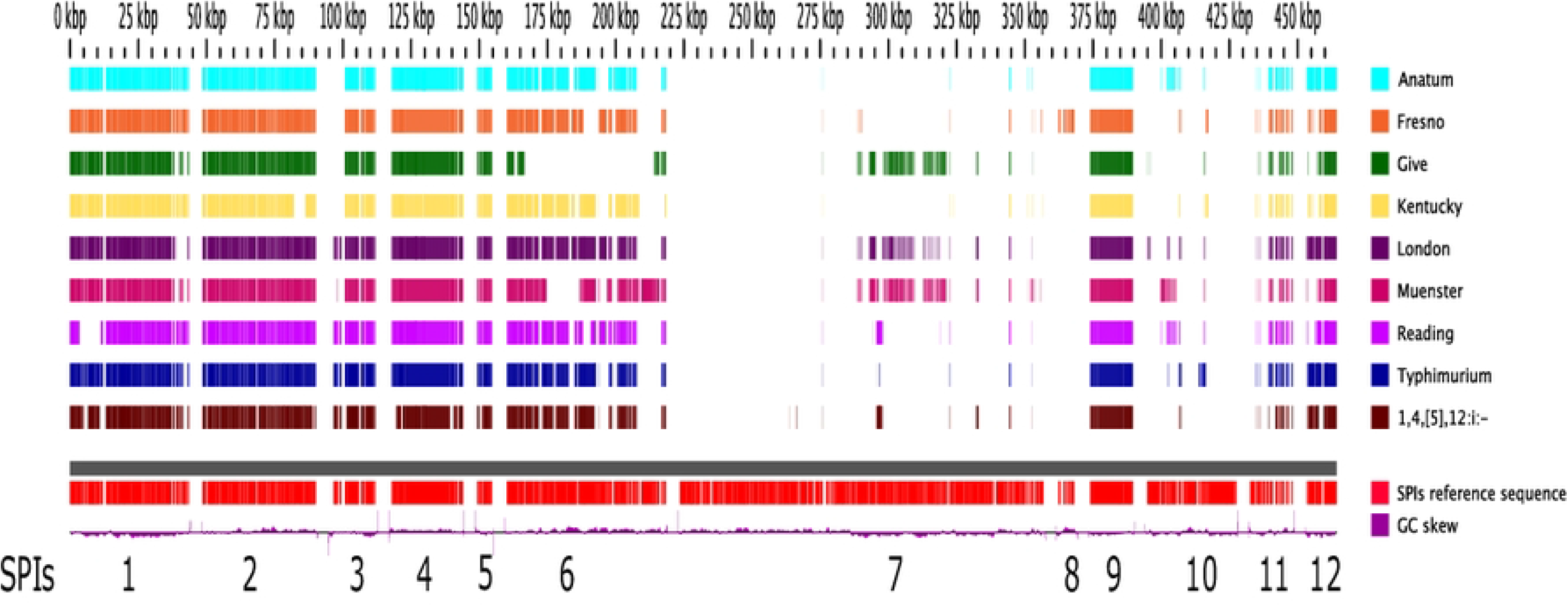
BLAST atlas analysis of 12 *Salmonella* pathogenicity islands (SPIs). The gray slot corresponds to the backbone. Above the backbone, each slot corresponds to a representative isolate of each serovar, whereas the slots below correspond to the reference sequence of each SPI (red) and the GC skew (purple). The numbers at the bottom indicate the SPI to which each fragment of the reference sequence belongs.

Overall, SPIs 1-5, 9, and 12 were rather conserved. We detected some partial deletions with different patterns across serovars close to the 3’ region of SPI-1 and the 5’ regions of SPIs 3, 11, and 12. The greatest variability occurred in SPI-6, which had four different versions across serovars, whereas it was not present in the serovar Give isolate. Moreover, most of the SPI-11 sequence was present in isolates of all serovars. However, as mentioned before, the *cdtB* islet, which encodes three key proteins (PltA, PltB, and CdtB) that form the so-called “typhoid toxin” [28], was only detected in isolates of serovars Give and Muenster.

Finally, analysis of the SNP clusters at the NCBI Pathogen Detection website revealed that isolates of all serovars, except Fresno, are closely related to strains involved in human infections. For instance, among the 93 isolates that formed the SNP cluster PDS000027247.46, one of our Anatum isolates (SRS5153609) was very close to several clinical strains from the USA. In particular, it was just 10 SNPs away from the clinical isolate PNUSA330427 (Fig 4). Likewise, one of our Typhimurium isolates (SRS5153603) was 2-9 SNPs away from other clinical isolates from the USA, Ireland, and the United Kingdom (Fig 4). Similar results were observed in the SNP clusters containing our isolates of serovars Reading (PDS000027459.15), London (PDS000027247.46), Kentucky (PDS000032811.10), Give (PDS000078457.2), and singleton monophasic 1,4,[5],12:i:- (PDS000176795.16).

**Figure 4.**
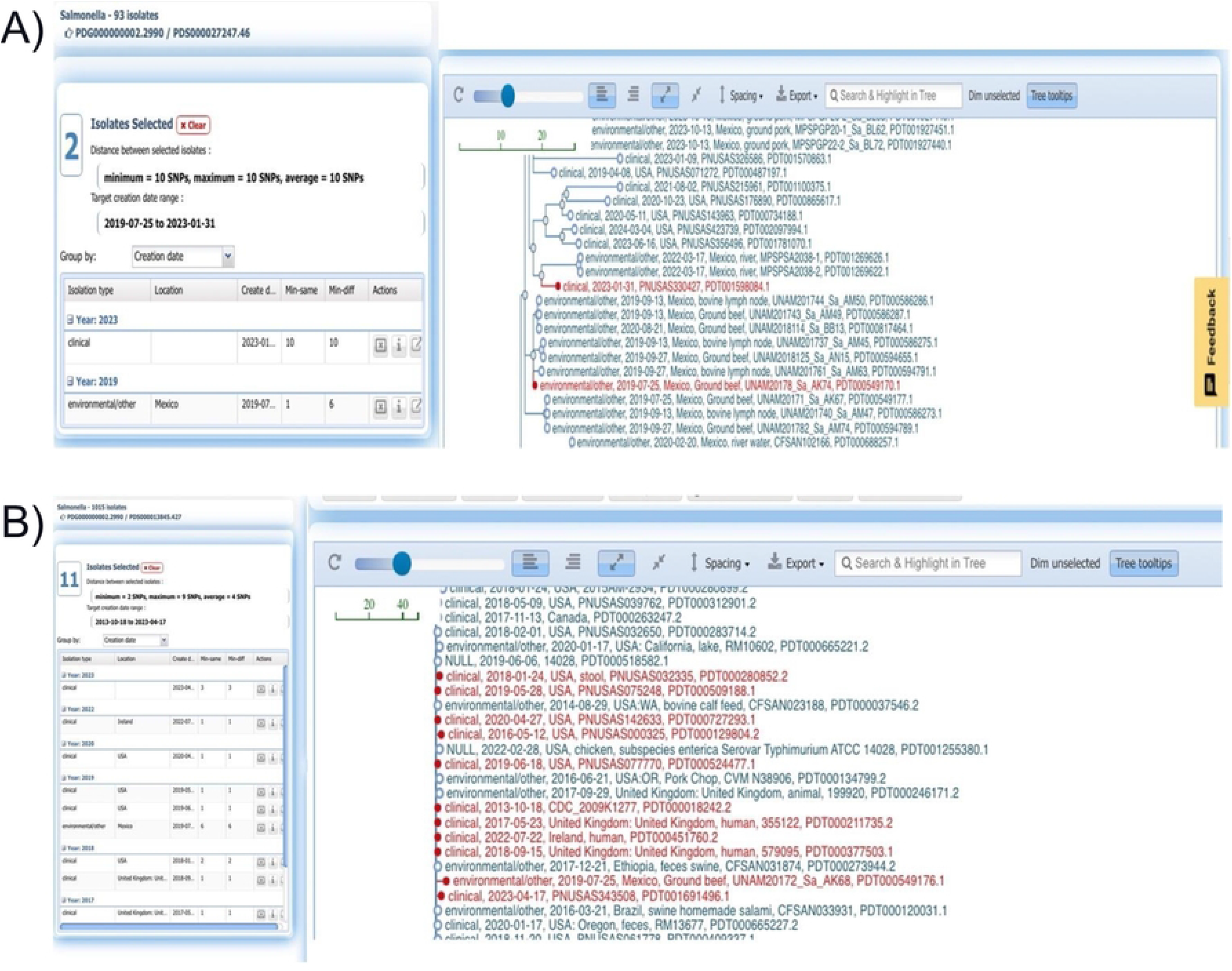
Fragments of two NCBI SNP clusters showing close genetic relatedness between our study isolates and clinical strains. A) SNP cluster PDS000027247.46 highlighting one of our Anatum isolates (UNAM20178_Sa_AK74) and a clinical strain from the USA (PNUSA330427). B) SNP cluster PDS000013845.427 highlighting one of our Typhimurium isolates (UNAM20172_Sa_AK68) and other clinical isolates from the USA, Ireland, and the United Kingdom.

## Discussion

In this study, we observed clonality between isolates from ground beef and lymph nodes (Fig 1), although both sample types were collected and analyzed separately. Moreover, character state reconstruction analysis documented that lymph nodes are the most likely source of the ancestors of isolates circulating in ground beef.

This pattern of genetic relatedness and evolutionary dynamics raises two relevant questions: 1) If the lymph nodes were not ground with the meat, why are some ground beef isolates still clonal with those from the lymph nodes? and 2) Is it possible that *Salmonella* circulating in the lymph nodes and ground beef are both of fecal origin?

One plausible explanation is that in carrier animals, some *Salmonella* circulating in the gut may reach the lymphatic system, creating a dual reservoir. Therefore, it may be incorporated into ground beef either via fecal contamination of the carcass or through the addition of lymph nodes during the grinding process.

Our results provide further evidence of the epidemiological role of lymph nodes in this regard. However, we could not assess that of the feces since our study was conducted at retail. Nonetheless, this hypothesis is consistent with recent research using a mouse model [29]. These authors documented that *Salmonella* migrates from the intestines to the lymph, either carried by dendritic cells or autonomously, and is captured by resident macrophages upon reaching the lymph nodes.

In cattle, the pathogen captured within macrophages has been shown to activate the immune response, which leads to its elimination [30]. However, studies with experimentally infected animals showed that it takes nearly one month for its total clearance [31]. Within this time frame, carrier animals could be sent for slaughter, increasing the risk of *Salmonella* dissemination along the food chain.

Given these facts, it is reasonable to speculate that *Salmonella* circulating in lymph nodes originates, at least in part, from the gut. This rationality is consistent with previous studies reporting indistinguishable PFGE subtypes between isolates from the feces and lymph nodes of healthy cattle [11, 12]. However, it has been difficult to infect lymph nodes through oral challenge without using high *Salmonella* concentrations [32], granting further research in this area.

Our results also document the ability of certain strains to persist within cattle populations, as shown by the close genetic relatedness of isolates of the same serovar, regardless of their collection year. These results agree with previous observations supporting vertical transmission (from a dam to her fetus) [33], as well as fecal oral transmission in cattle [34]. They are also in line with the documented ability of *Salmonella* to establish long-term (up to one year) dormant infections in the mesenteric lymph nodes of mice [35].

Along the same lines, it has been observed that the frequency of lymph nodes carrying *Salmonella* increases as feedlot cattle move into the later stages of the fattening period [36]. However, further research is needed since the factors associated with this transmission dynamics, or the mechanisms allowing the pathogen to persist for extended periods across cattle cohorts, have not been identified.

Results of the virulence and pathogenicity islands profile were consistent with those of the phylogenetic analysis. Most variation occurred between serovars. For instance, *Salmonella* virulence plasmid genes were present only in isolates of serovar Typhimurium, which were collected from ground beef but not from lymph nodes. *Salmonella* strains carrying the virulence plasmid are known to exhibit hypervirulent phenotypes, which is inconsistent with their isolation from apparently healthy animals [37]. Perhaps, as previously suggested by other researchers [38], under scenarios favoring a continuous fecal-oral infection cycle, *Salmonella* may successfully replicate without entering the intestinal epithelium. Therefore, the pathogen may not express the genes involved in acute salmonellosis during persistent infections settings [39].

Other minor variations in virulence genes involved factors related to adherence and iron uptake and metabolism. However, *Salmonella* has functional redundancy in these pathogenic processes. Hence, it is unlikely that these differences are related to lymph node specialization since predominant serovars (i.e. Anatum, Reading, London) had a rather uniform virulence profile, regardless of their isolation source. Furthermore, the limited number of isolates carrying genes encoding the typhoid toxin and hemolysin E is consistent with their narrow distribution among non-typhoidal *Salmonella* [40].

Regarding pathogenicity islands, different versions occurred between serovars. In general, the rather uniform pathogenicity islands profile did not support an association between the isolation source and the virulence repertoire of our study isolates. Moreover, the high conservation of SPIs 1-5 suggests that all these isolates have the essential factors to colonize mammals. In fact, previous studies have demonstrated that even the disruption of several genes of SPIs 1-3 does not affect the ability of *Salmonella* strains to colonize cattle, pigs, and chicken [41].

Taken together, the results of the virulence and pathogenicity islands indicate that *Salmonella* strains circulating in apparently healthy animals have a conserved virulence repertoire. Hence, the establishment of the carrier state in cattle depends on factors other than the differences in the major virulence genes and pathogenicity islands considered in this study.

These findings were further supported by analysis of the phylogenetic tree at the NCBI Pathogen Detection website. It showed that isolates of all serovars except Fresno were genetically close to strains involved in human infections in different countries. Hence, there is no doubt that isolates circulating in apparently healthy animals pose a risk to public health.

Overall, this study confirms the role of lymph nodes as a key reservoir of *Salmonella* circulating in ground beef. It also showed that carrier animals are critical for disseminating the pathogen along the food continuum, although we did not identify changes in the virulence repertoire that could be associated with attenuated virulence or the establishment of the carrier state in cattle. In general, our study isolates had a similar virulence and pathogenicity islands profile, regardless of their isolation source, whereas strains of most serovars were genetically close to those involved in human infections in other countries. Further research is required to decipher the mechanisms by which *Salmonella* causes subclinical infections in cattle and persists for long periods in farm environments.

## References

1. CDC. National Enteric Disease Surveillance: Salmonella Annual Report, 2010: US Department of Health and Human Services; 2013 [cited 2024 April 3]. Available from: http://www.cdc.gov/ncezid/dfwed/pdfs/salmonella-annual-report-appendices-2010-508c.pdf.

2. CDC. Outbreak of Salmonella infections linked to ground beef: Centers for Disease Control and Prevention (CDC) 2018 [cited 2024 April 3]. Available from: https://archive.cdc.gov/#/details?url=https://www.cdc.gov/salmonella/newport-10-18/index.html.

3. Organization WH. WHO estimates of the global burden of foodborne diseases 2015 [cited 2024 April 3]. Available from: https://www.who.int/publications/i/item/9789241565165.

4. Webb HE, Brichta-Harhay DM, Brashears MM, Nightingale KK, Arthur TM, Bosilevac JM, et al. Salmonella in peripheral lymph nodes of healthy cattle at slaughter. Frontiers in Microbiology. 2017;8: 2214. doi: 10.3389/fmicb.2017.02214. PMID: 29170662

5. Ayala D, Nightingale K, Narvaez-Bravo C, Brashears MM. Molecular characterization of Salmonella from beef carcasses and fecal samples from an integrated feedlot and abattoir in Mexico. Journal of Food Protection. 2017;80(12): 1964–72. doi: 10.4315/0362-028x.jfp-17-157. PMID: 29130766

6. Delgado-Suárez EJ, Selem-Mojica N, Ortiz-López R, Gebreyes WA, Allard MW, Barona-Gómez F, Rubio-Lozano MS. Whole genome sequencing reveals widespread distribution of typhoidal toxin genes and VirB/D4 plasmids in bovine-associated nontyphoidal Salmonella. Scientific Reports. 2018;8(1): 9864. doi: 10.1038/s41598-018-28169-4.

7. Rodriguez-Rivera L, Moreno Switt AI, Degoricija L, Fang R, Cummings CA, Furtado MR, et al. Genomic characterization of Salmonella Cerro ST367, an emerging Salmonella subtype in cattle in the United States. BMC Genomics. 2014;15: 427. doi: 10.1186/1471-2164-15-427.

8. Locke SR, Pempek JA, Meyer R, Portillo-Gonzalez R, Sockett D, Aulik N, Habing G. Prevalence and sources of Salmonella lymph node infection in special-fed veal calves. Journal of Food Protection. 2022;85(6): 906–17. doi: 10.4315/JFP-21-410. PMID: 35146524

9. Palós Gutiérrez T, Rubio Lozano MS, Delgado Suárez EJ, Rosi Guzmán N, Soberanis Ramos O, Hernández Pérez CF, Méndez Medina RD. Lymph nodes and ground beef as public health importance reservoirs of Salmonella spp. Revista Mexicana de Ciencias Pecuarias. 2020;11(3): 795–810. doi: 10.22319/rmcp.v11i3.5516.

10. Nickelson KJ, Taylor TM, Griffin DB, Savell JW, Gehring KB, Arnold AN. Assessment of Salmonella prevalence in lymph nodes of U.S. and Mexican cattle presented for slaughter in Texas. Journal of Food Protection. 2019;82(2): 310–5. doi: 10.4315/0362-028X.JFP-18-288. PMID: 30682264

11. Gragg SE, Loneragan GH, Nightingale KK, Brichta-Harhay DM, Ruiz H, Elder JR, et al. Substantial within-animal diversity of Salmonella isolates from lymph nodes, feces, and hides of cattle at slaughter. Applied and Environmental Microbiology. 2013;79(15): 4744–50. doi: 10.1128/AEM.01020-13.

12. Munoz-Vargas L, Finney SK, Hutchinson H, Masterson MA, Habing G. Impact of clinical salmonellosis in veal calves on the recovery of Salmonella in lymph nodes at harvest. Foodborne Pathogens and Disease. 2017;14(11): 678–85. doi: 10.1089/fpd.2017.2303.

13. Andrews S. FastQC: a quality control tool for high throughput sequence data 2010 [cited 2024 April 3]. Available from: http://www.bioinformatics.babraham.ac.uk/projects/fastqc.

14. Bolger AM, Lohse M, Usadel B. Trimmomatic: A flexible trimmer for Illumina sequence data. Bioinformatics. 2014;15: 2114–20. doi: 10.1093/bioinformatics/btu170.

15. Bankevich A, Nurk S, Antipov D, Gurevich AA, Dvorkin M, Kulikov AS, et al. SPAdes: a new genome assembly algorithm and its applications to single-cell sequencing. Journal of Computational Biology. 2012;19(5): 455–77. doi: 10.1089/cmb.2012.0021. PMID: 22506599

16. Gurevich A, Saveliev V, Vyahhi N, Tesler G. QUAST: quality assessment tool for genome assemblies. Bioinformatics. 2013;29(8): 1072–5. doi: 10.1093/bioinformatics/btt086. PMID: 23422339

17. Delgado-Suarez EJ, Palos-Guiterrez T, Ruiz-Lopez FA, Hernandez Perez CF, Ballesteros-Nova NE, Soberanis-Ramos O, et al. Genomic surveillance of antimicrobial resistance shows cattle and poultry are a moderate source of multidrug resistant non-typhoidal Salmonella in Mexico. PLoS One. 2021;16(5): e0243681. doi: 10.1371/journal.pone.0243681. PMID: 33951039

18. Brettin T, Davis JJ, Disz T, Edwards RA, Gerdes S, Olsen GJ, et al. RASTtk: A modular and extensible implementation of the RAST algorithm for building custom annotation pipelines and annotating batches of genomes. Scientific Reports. 2015;5(8365). doi: 10.1038/srep08365.

19. Kaas RS, Leekitcharoenphon P, Aarestrup FM, Lund O. Solving the problem of comparing whole bacterial genomes across different sequencing platforms. PLoS One. 2014;9(8): e104984. doi: 10.1371/journal.pone.0104984. PMID: 25110940

20. Stamatakis A, Hoover P, Rougemont J. A rapid bootstrap algorithm for the RAxML web servers. Systematic Biology. 2008;57(5): 758–71. doi: 10.1080/10635150802429642. PMID: 18853362

21. Miller MA, Pfeiffer W, Schwartz T. Creating the CIPRES Science Gateway for inference of large phylogenetic trees. Proceedings of the Gateway Computing Environments Workshop (GCE), 14 Nov 2010, New Orleans, LA pp 1–8. 2010.

22. Letunic I, Bork P. Interactive tree of life (iTOL) v5: an online tool for phylogenetic tree display and annotation. Nucleic Acids Research. 2021;49(W1): W293–W6. doi: 10.1093/nar/gkab301. PMID: 33885785

23. Madisson WP, Madisson DR. Mesquite: A modular system for evolutionary analysis 2021 [cited 2024 April 3]. Available from: http://www.mesquiteproject.org.

24. Chen L, Zheng D, Liu B, Yang J, Jin Q. VFDB 2016: hierarchical and refined dataset for big data analysis--10 years on. Nucleic Acids Research. 2016;44(D1): D694–7. doi: 10.1093/nar/gkv1239. PMID: 26578559

25. Yoon SH, Park YK, Kim JF. PAIDB v2.0: Exploration and analysis of pathogenicity and resistance islands. Nucleic Acids Research. 2015;43(Database issue): D624–30. doi: 10.1093/nar/gku985. PMID: 25336619

26. Petkau A, Stuart-Edwards M, Stothard P, Van Domselaar G. Interactive microbial genome visualization with GView. Bioinformatics. 2010;26(24): 3125–6. doi: 10.1093/bioinformatics/btq588. PMID: 20956244

27. Herrero-Fresno A, Olsen JE. Salmonella Typhimurium metabolism affects virulence in the host - A mini-review. Food Microbiology. 2018;71: 98–110. doi: 10.1016/j.fm.2017.04.016. PMID: 29366476

28. Spano S, Ugalde JE, Galan JE. Delivery of a Salmonella Typhi exotoxin from a host intracellular compartment. Cell Host Microbe. 2008;3(1): 30–8. doi: 10.1016/j.chom.2007.11.001. PMID: 18191792

29. Bravo-Blas A, Utriainen L, Clay SL, Kastele V, Cerovic V, Cunningham AF, et al. Salmonella enterica serovar Typhimurium travels to mesenteric lymph nodes both with host cells and autonomously. The Journal of Immunology. 2019;202(1): 260–7. doi: 10.4049/jimmunol.1701254. PMID: 30487173

30. Arsenault RJ, Brown TR, Edrington TS, Nisbet DJ. Kinome analysis of cattle peripheral lymph nodes to elucidate differential response to Salmonella spp. Microorganisms. 2022;10(1). doi: 10.3390/microorganisms10010120. PMID: 35056570

31. Edrington TS, Loneragan GH, Genovese K, Hanson DL, Nisbet DJ. Salmonella persistence within the peripheral lymph nodes of cattle following experimental Inoculation. Journal of Food Protection. 2016;79(6): 1032–5. doi: 10.4315/0362-028X.JFP-15-325.

32. Brown TR, Edrington TS, Genovese KJ, Loneragan GH, Hanson DL, Nisbet DJ. Oral Salmonella challenge and subsequent uptake by the peripheral lymph nodes in calves. Journal of Food Protection. 2015;78(3): 573–8. doi: 10.4315/0362-028X.JFP-14-416. PMID: 25719883

33. Hanson DL, Loneragan GH, Brown TR, Nisbet DJ, Hume ME, Edrington TS. Evidence supporting vertical transmission of Salmonella in dairy cattle. Epidemiology and Infection. 2016;144(5): 962–7. doi: 10.1017/S0950268815002241. PMID: 26419321

34. Dodd CC, Renter DG, Shi X, Alam MJ, Nagaraja TG, Sanderson MW. Prevalence and persistence of Salmonella in cohorts of feedlot cattle. Foodborne Pathogens and Disease. 2011;8(7): 781–9. doi: 10.1089=fpd.2010.0777.

35. Li Q. Mechanisms for the invasion and dissemination of Salmonella. Canadian Journal of Infectious Diseases and Medical Microbiology. 2022;2022: 2655801. doi: 10.1155/2022/2655801. PMID: 35722038

36. Belk AD, Arnold AN, Sawyer JE, Griffin DB, Taylor TM, Savell JW, Gehring KB. Comparison of Salmonella prevalence rates in bovine lymph nodes across feeding stages. Journal of Food Protection. 2018;81(4): 549–53. doi: 10.4315/0362-028X.JFP-17-254. PMID: 29513102

37. Guiney DG, Fierer J. The role of the spv genes in Salmonella pathogenesis. Frontiers in Microbiology. 2011;14(2): 129. doi: 10.3389/fmicb.2011.00129. PMID: 21716657

38. Ginocchio CC, Rahn K, Clarke RC, Galán JE. Naturally occurring deletions in the centisome 63 pathogenicity island of environmental isolates of Salmonella spp. Infection and Immunity. 1997;65(4): 1267–72. doi: 10.1128/iai.65.4.1267-1272.1997.

39. Barat S, Steeb B, Mazé A, Bumann D. Extensive in vivo resilience of persistent Salmonella. PLoS One. 2012;7(7): e42007. doi: 10.1371/journal.pone.0042007.

40. Figueiredo R, Card R, Nunes C, AbuOun M, Bagnall MC, Nunez J, et al. Virulence characterization of Salmonella enterica by a new microarray: Detection and evaluation of the cytolethal distending toxin gene activity in the unusual host S. Typhimurium. PLoS One. 2015;10(8): e0135010. doi: 10.1371/journal.pone.0135010. PMID: 26244504

41. Chaudhuri RR, Morgan E, Peters SE, Pleasance SJ, Hudson DL, Davies HM, et al. Comprehensive assignment of roles for Salmonella Typhimurium genes in intestinal colonization of food-producing animals. PLoS Genetics. 2013;9(4): e1003456. doi: 10.1371/journal.pgen.1003456. PMID: 23637626

